# On Biases of Attention in Scientific Discovery

**DOI:** 10.1101/2020.04.08.002378

**Authors:** Uriel Singer, Kira Radinsky, Eric Horvitz

## Abstract

How do nuances of scientists’ attention influence what they discover? We pursue an understanding of the influences of patterns of attention on discovery with a case study about confirmations of protein-protein interactions over time. We find that modeling and accounting for attention can help us to recognize and interpret biases in databases of confirmed interactions and to better understand missing data and unknowns in our fund of knowledge.

## Introduction

Indian Philosopher Jiddu Krishnamurthy said that “the finding is not in the future–it is there, where you do not look.” In the spirit of Krishnamuthy’s reflection, we investigate how patterns of attention in scientific discovery can influence the state of knowledge in a discipline in ways that are not recognized. We show with a case study how biases of attention can influence the distribution of hypotheses that are proposed and confirmed over time. These biases can lead to implicit assumptions about the significance of hypotheses that remain unconfirmed or unstated. We focus on the illustrative example of the growing fund of knowledge about interactions among proteins. We find evidence via an analysis of snapshots over time of known protein-protein interactions (PPI) that discoveries about protein interactions exhibit a bias rooted in scientists’ attention on recent discoveries. Such biases may be based in several factors, including a sequencing of attention on specific sets of biochemical pathways of interest, and a growth of understanding of these systems via testing hypotheses about the role of proteins with one or more proteins already known to be interacting with one another in the systems.

Decades of research have yielded large databases of protein-protein interactions. These databases have played a critical role in biomedicine, enabling the construction of biochemical cascades and larger protein interaction networks. The interaction data and resulting representations of metabolic, structural, and regulatory processes have been valuable in understanding the etiology of diseases and in identifying promising therapies, including efforts at prioritizing pharmacological targets.

The human genome codes for approximately 30,000 proteins ^1^. Given two proteins *p*_*i*_, *p*_*j*_, we can represent the probability of observing a confirmation of an interaction between them over some period of time, *pr*(*p*_*i*_ ↔ *p*_*j*_) as a chain of two probabilities: (1) the probability of an experiment being performed to check the interaction during that period, and, (2) the probability that the proteins will be found to interact, given that the experiment is carried out. Thus, taking both probabilities into consideration, the likelihood of seeing an interaction confirmed is:

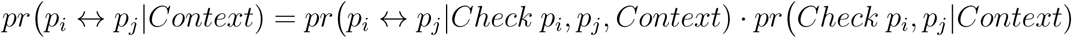

where we condition all probabilities on the scientific context, which includes background knowledge about protein interactions.

We focus on the probability that a protein-protein interaction is checked experimentally, *pr*(*Check p*_*i*_, *p*_*j*_|*Context*). This probability can be further decomposed into two probabilities: (1) the probability that scientists are interested in performing a specific experiment—their *attention* to seeking confirmation or disconfirmation of the hypothesis and (2) the probability that they have the required scientific tools, including experimental resources and affordances, given interest in doing a study, written as follows:

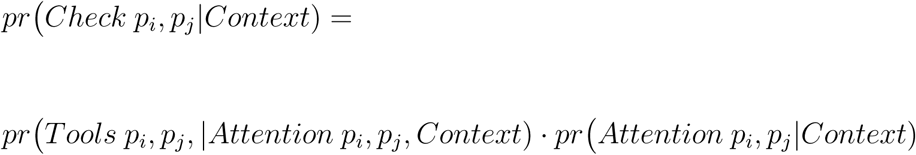

Breaking out these probabilities can provide a valuable perspective on the current state of knowledge, the influence of enabling tools, and on biases of attention in different areas of science. We consider the growth of the database of protein-protein interactions as a source of insights about the influences of attention in scientific discovery. Findings about systematic biases introduced by attentional considerations on knowledge about protein interactions may generalize to other scientific pursuits.

We find that *pr*(*Check p*_*i*_, *p*_*j*_|*Context*) shows patterns of biases of attention, and that attentional influences on the choice of pairs of proteins that are checked for interactions lead in turn to biases in the newly confirmed interactions that are added to the growing PPI database. Biases of attention often lurk as implicit because their role is not overtly considered in interpretations of confirmed, disconfirmed, and unknown protein interactions. Such implicit biases may fuel systematic patterns of unknowns in scientific databases. To visualize the influence of attention, we consider the database of all known protein interactions at the end of each calendar year for a span of years under consideration. For each year, we represent the protein-interaction database as a graph, where proteins are nodes and edges represent interactions among proteins that are confirmed by the end of the respective year. Within any annual PPI graph, we define the *protein distance* between any two proteins as the length of the shortest chain connecting the proteins in the graph (see Method “Data” for more details on the database creation).

We have found that new discoveries about protein interactions in a following year are highly skewed towards protein interactions with small distances in the PPI graph for the current year, as captured in Figure 2. In the graph one can see the distribution of discovered edge distances a year before discovery normalized by the distribution of all possible edge distances, creating a probability graph. We had first discovered this phenomena during studies exploring the use of machine learning to predict future interactions. During these investigations, we were surprised and intrigued by the dominating evidential strength of the proximity of proteins in the PPI graph for predicting future confirmations.

**Figure 1:**
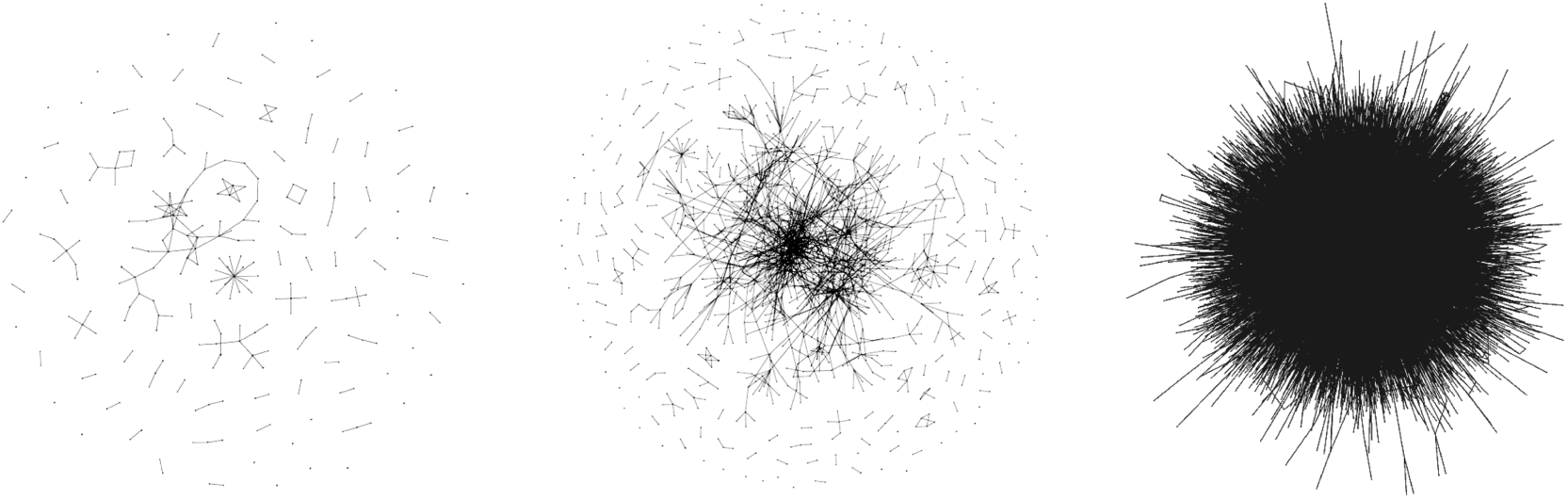
Protein-protein interactions represented as a graph where nodes are proteins and edges are interactions. From left to right, expansion of the graph over the years.

**Figure 2:**
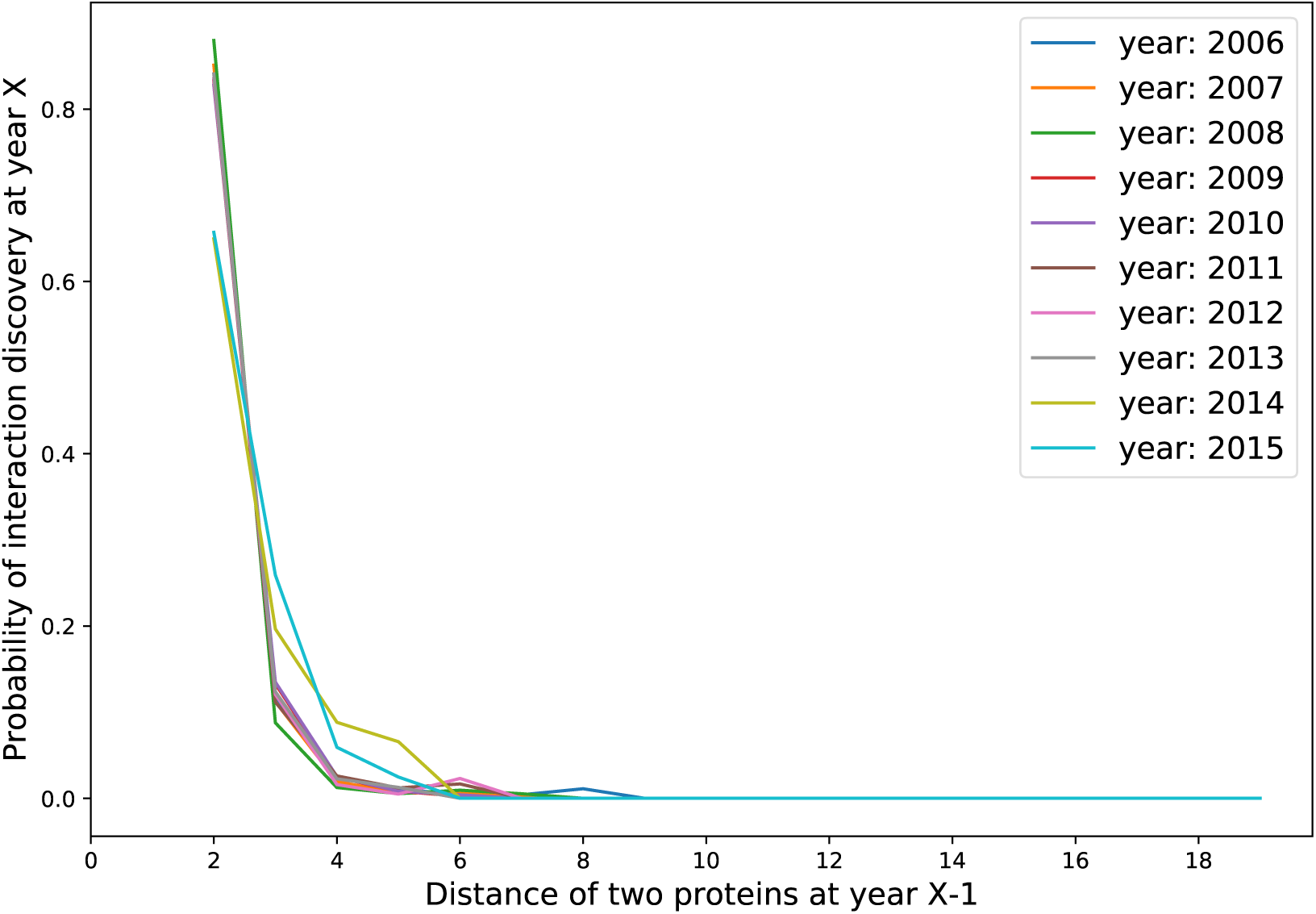
Probability that a protein interaction will be confirmed in the next year as a function of the distance among two proteins in the interaction graph in a current year. Each color represents a different current year.

We could see no biological rationale to explain the large observed skew of discoveries based on adjacencies. We hypothesized that the observation are based in biases in the attention of scientists to continuing exploration of proteins and processes that have been most recently studied. In the absence of attentional influences, we would expect a more uniform distribution in distances among proteins that are found to interact in the next year. We set out to probe the presence and influence of these potential biases on the sequencing of discoveries about protein interactions given the likely presence of similar biases of attention in other areas of science. We believe that unearthing and characterizing such biases in research on protein interactions and in other scientific endeavors will be important for interpreting distributions of knowns and unknowns at different points in time in the maturation of fields.

### On Countering Attentional Influences

We now pursue means for recognizing and countering the influence of biases of attention in discoveries about protein interactions. We study the effect of proximity in the PPI graph on the inferred probability of future discoveries via the construction of statistical models that predict that untested pairs of proteins will be found to interact–should they be studied. We explore the ability of predictive models to forecast future discoveries when they are trained on data about protein interactions known at an earlier date. We then consider the power of the use of representations of the data that can be viewed as less biased in that they do not rely on graph proximity.

By definition, proteins at distance *d* = 1 are already confirmed to interact. We seek to compare the growth in knowledge about the interactions between proteins at *d* = 2 versus proteins that are separated by greater distances in the graph. This comparison can reveal the potential for an alternate sequencing of discoveries about protein interactions in a world without these biases. We perform a counterfactual analysis based on matching ^2^ (see Method “Causal Inference and matched sets” for more details). We identify matched proteins via two approaches. In the first approach, we consider protein biological features by applying an embedding technique on its amino acid code inspired by ^3^. By concatenating three types of descriptors: composition (C), transition (T), and distribution (D), we end up with a feature vector for each protein, while the edge feature vector is calculated as the average of its two protein biological feature vectors. In a second approach, we match protein pairs by using graph embeddings (see Method “Node2vec” for more details) via the use of neural graph learning methods.

Based on the graph embeddings, we train a logistic regression model on pairs of proteins in the PPI graph from the year 2012. We use the trained model to infer the probability that an edge will appear in the period of 2013–2015. We compare four sets of predictions:

1. *Biased*: Predictions considering proteins and their original edges in the PPI graph as features.
2. *Debiased, graph embedding*: Prediction on edges matched via graph embeddings with ‘co-sine’ distance metric. That is, the matching is applied between protein pairs with edges at *d* > 2 to the proteins with edges at *d* = 2 by using their graph embeddings similarity.
3. *Debiased, amino acid*: Prediction on edges matched between edges at *d* > 2 and edges at *d* = 2 based on their similarity based on biological features with ‘canberra’ distance metric.
4. *Random*: Prediction on the matched edges, if they will become edges. Matching is applied between the original edges at *d* > 2 to edges at *d* = 2 by random sampling. This random sampling is interesting to neglect false obvious insights.

We use as training data protein interactions in the PPI graph from the year 2012 and use the learned model to perform predictions about interactions discovered in the period of 2013–2015. We sample random false edges and report as a metric of performance the area under the receiver-operator characteristic curve (AUC) of the predictive model on these later discoveries of interactions for the four different sets of predictions.

In Table 1, we show for each distance the AUC for each type of prediction method in predicting the interactions discovered in 2013–2015. It is evident that predicting using graph-based matching significantly improves the predictions of new discoveries, per measures of the boost in AUC. We note that, in the most recent years being studied, 2014-15, interactions appear to be discovered via more advanced techniques and tools, leading the test set to be less attentionally biased than in earlier years. Practically, we train the predictive model on a biased training set and predict on a less biased test set. We attempt to weaken biases of attention by applying graph-based matching than using the given ‘biased’ edges of 2012. The new techniques still hold bias, indicating that the graph-based matching of today’s known PPI, could lead to even better predictions. In other words, (*Check p*_*i*_, *p*_*j*_|*Context*) in later years gets less dependent on the proximity of *i* and *j*, and therefore *pr*(*p*_*i*_ ↔ *p*_*j*_|*Context*) is less biased.

**Table 1:** AUC results over the given distances for each method

| distance | Biased | Debiased, graph embedding | Debiased, amino acid | Random |
| --- | --- | --- | --- | --- |
| 2 | 0.788 | 0.788 | 0.788 | 0.499 |
| 3 | 0.623 | **0.656** | 0.638 | 0.493 |
| 4 | 0.571 | **0.635** | 0.566 | 0.514 |
| 5 ≤ | 0.518 | **0.626** | 0.587 | 0.490 |
| *all* | 0.726 | **0.770** | 0.719 | 0.498 |

While our focus has been elucidating attentional infuences on sequence of discoveries about protein interactions, we foresee that such predictive models could help provide scientists with guidance on exploratory directions by helping scientists to see around attentional biases of proximity.

### Defining Bias of Attention

We can further define attentional bias. By using the *graph-debias* method from Section “On Countering Attentional Influences” we can compare the individual edge scores between the original and matched edges. The *graph-debias* method is produced by identifying pairs of proteins drawn from all pairs at each distance *d* = *n* to their closest matches at distance *d* = 2. The distance metric is applied on the edge embedding which is simply defined by the mean of the two nodes’ embeddings. After performing the matching, we calculate an individual treatment effect (ITE), which is the difference of an edge score for the 2013–2015 period of the original *d* = 2 edges and the matched edges: *y*_*i*_(*y*_2(*i*)_ − *y*_*n*(*i*)_), where each sample *i* is a potential edge, *y*_*n*(*i*)_ is the original edge probability being discovered in later years, *y*_2(*i*)_ is the matched edge probability, and *y*_*i*_ is the true label of the original edge, where *y*_*i*_ = 1 if an edge forms between the proteins by the end of year 2015, and *y*_*i*_ = −1 if not.

The value of *y*_*i*_(*y*_2(*i*)_ − *y*_*n*(*i*)_) can be between −1 and 1, where −1 is assigned if an edge is predicted correctly between the original edge at *d* > *n* but incorrectly for the matched edge at *d* = 2, 1 if an edge is predicted correctly between the matched edge but is not for the original edge, and 0 if there’s no difference on the edge prediction for the original and matched edges.

We then calculate the average treatment effect (ATE) for each distance:

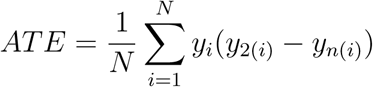

Observing *ATE* > 0 means that the matched edges are better in predicting edges than the original edges. The closer the value of *ATE* is to 1, the better the predictions of the matched edges over the original edges. In the same way, the closer the value of *ATE* is to −1, the worse the predictions of the matched edges will be over the original edges. We propose that the average treatment effect provides a valuable characterization of bias in attention for discovery of protein interactions.

As displayed in Figure 3, we can see that the edges for the matched edges perform better than the original edges at *d* > *n*. We observe that the larger the distance, the larger the effect of moving from the original to the matched edges.

**Figure 3:**
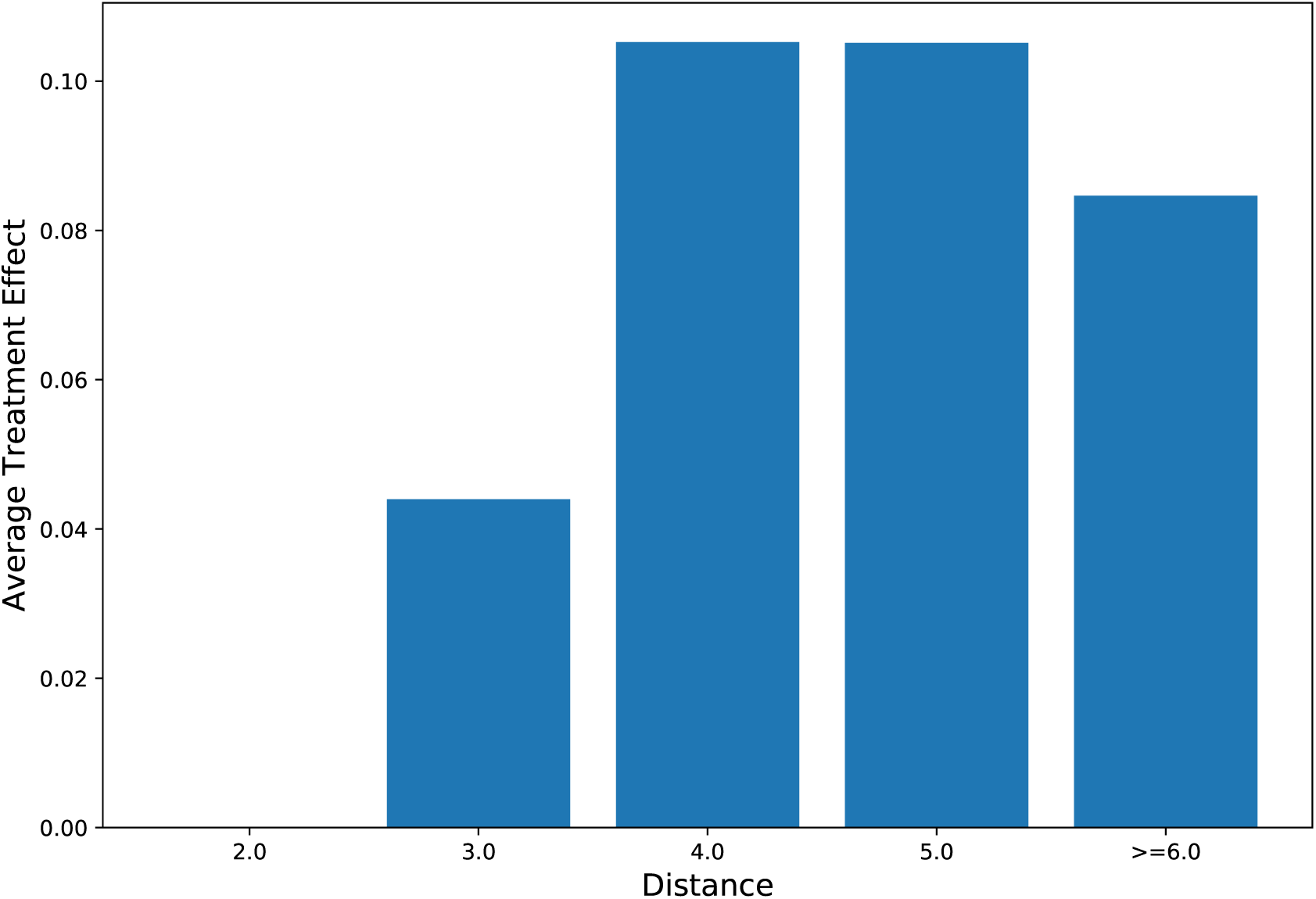
Average treatment effect as a function of the distance of matched edges.

### Looking to the Future

We examined the sequence of confirmations about protein-protein interactions over time and found evidence of biases of locality of scientists’ focus of attention: scientists weight the formulation and confirmation of hypotheses toward previously confirmed protein-protein interactions. This attentional bias has shaped the progression of knowledge about protein interactions. Similar attentional factors may play an important role across the sciences. Biases of discovery based on temporal and conceptual proximity highlight the potential value of adding to research portfolios the formulation and pursuit of hypotheses that make more distant leaps in conceptual spaces. The findings for protein-protein interactions emphasize our need to better understand biases of attention across the sciences so as to broaden our understandings of the structure and implications of unexplored hypotheses and missing data. We can also work to counter potential biases of locality by nurturing research environments that promote exploration of more distant conceptual relationships and of making serendipitous discoveries ^4^ that jump beyond the frontier of current knowledge.

## Methods

### Data

The protein interaction information used in the study was drawn from HINTdb ^5^. HINTdb contains a list of all protein interactions, where each interaction includes a list of all articles mentioning it. We perform a preprocessing phase, where we identify the discovery date of an interaction as the earliest associated article publication date.

Given a date, we create a graph that includes all interactions with their associated discovery dates, earlier than the given date. By using this method, we receive a graph that evolves over time. Protein interactions with articles or articles with dates were not included in the graph. In the experiments, the PPI graph of year *y* includes all interactions among proteins that are confirmed by the end of year *y*.

### Node2vec

We opted to work with node2vec ^6^, which has achieved state-of-the-art performance on multiple benchmarks. Node2vec is based on word2vec ^7^, which is a framework for feature learning representation of words. The framework receives as input a text corpus and outputs an embedding in a low-dimensional space of size *d* (a hyperparameter) for each word. By trying to predict the words’ neighbors, word2vec is able to create for each word an embedding that represents the words’ semantic meaning. Node2vec generalizes word2vec for the graph domain, where intuitively, each node is regarded as a word. The algorithm creates the equivalent to sentences by performing random graph walks starting from all nodes (i.e., each node sampled in the random walk is a word in the sentence).^1^

### Causal Inference and matched sets

We use the health domain in order to explain the method in an intuitive way: Assume that we have two groups of patients, where one group has received a treatment and the other has not. We know the outcomes of all the patients, for example their blood pressure. Being able to answer the question of what would be the treatment effect on a specific patient or maybe in average is a great deal that causal inference techniques can provide. A reduction from the health domain to the PPI domain can be done by looking at a potential edges as patients, at the treatments as the distances and at the outcomes as the answer will they be an edge in the future. In this way we can apply the same techniques on the PPI domain and answer the question raised in the beginning of the section. ^2^ showed the power of the matching, which means, given two groups that differ in the treatments they received, find for each individual in the first group its closest match in the second group (in some feature vector space). By finding these matches, we are able to find the individual treatment affect (ITE) by the difference of its own outcome and its matched outcome. In order to receive the average treatment effect (ATE) we simply average over all individuals. We can apply matching on the PPI domain by finding for each edge with distance *d* > 2 its match in some feature space with distance *d* = 2 and calculate straightforward the distance effect.

## Footnotes

1 We used the implementation publicly published by the authors: github.com/aditya-grover/node2vec

